# Plasma levels of Ceramides and their association with Hematocrit and Thrombocytopenia in Severe and Non-severe dengue

**DOI:** 10.1101/2022.06.22.497160

**Authors:** Aashika Raagavi Jeanpierre, Vignesh Mariappan, Siva Ranganathan Green, Srinivasa Rao Mutheneni, Shashikala P, Agieshkumar Balakrishna Pillai

## Abstract

**Background:** Plasma leakage due to endothelial permeability is regarded as a hallmark process in the pathophysiology of severe dengue. In recent years, the importance of ceramide in regulating vascular tone during viral infection and metabolic diseases has gained attention. The present study aimed to evaluate the plasma levels of ceramide and its association with plasma leakage in dengue patients.

**Methods:** The study involved 30 dengue samples involving severe dengue (SD-10), Dengue with Warning sign (DWW-10), Dengue without Warning Sign (DWOW-10), along with other febrile illness (OFI-10) controls. Samples were collected on the day of admission (DOA), day of defervescence (DOD), and day of convalescence (DOC). Total plasma ceramides (d18:1/22:0) were quantified using RP-HPLC. The correlation between ceramides and hematocrit/platelet count was evaluated using Spearman Rho Correlation. All the statistical analyses were performed using SPSS software.

**Results:** During the febrile phase, a significant (P≤0.05) decrease in the levels of ceramides was observed in dengue compared to OFI (control). Towards the defervescences, the ceramides levels were substantially (P ≤ 0.001) elevated in dengue groups compared to baseline. Most importantly, the ceramide levels were found to be higher in SD patients compared to non-severe dengue (DWW & DWOW) and OFI, particularly at the critical phase of infection. We observed a significant negative correlation (r = -0.867, P ≤0.001) between the platelet count and ceramide levels in SD subjects. Notably, a negative correlation was observed between ceramide and hematocrit during the defervescence phase (R = -0.355, P≤0.05) in the AD group.

**Conclusion:** Elevated levels of circulating plasma ceramides during the defervescence phase of severe dengue show an essential role of ceramides in disease pathogenesis, however further studies are required to ascertain it.

## Introduction

Dengue fever is a resurgent arbovirus disease that causes 100-400 million infections each year (1). Dengue disease is characterized by fever, headache, muscle, and joint pains, rash, nausea, and vomiting. Some infections result in dengue hemorrhagic fever (DHF), a syndrome that in its most severe form can threaten the patient’s life, primarily through increased vascular permeability and shock (2). Nevertheless, on the 3rd to 7th day of the illness, as the fever subsides, some patients manifest severe signs like endothelial dysfunction, which induces vascular permeability. Thrombocytopenia and vascular dysfunction are the cardinal features of dengue pathogenesis. Currently, there is no commercially available vaccine for dengue, but treatment is supportive. Early prediction of disease outcomes could provide prompt medical attention and reduce the risk of fatality (2)

The exact mechanism that regulates vascular dysfunction and platelet destruction in the pathogenesis of dengue remains obscure. Recently, we have reported the role of platelet lipids in the pathogenesis of dengue(3). Lipids play a role in structural, energy storage, and cellular barriers. The role of lipids in the pathogenesis of single-stranded RNA virus disease is not well established. We have reported an alteration in the platelet fatty acids in severe dengue cases(4). A lipidomic study on dengue plasma revealed a differential expression of phospholipids and sphingolipids in the severe form of dengue compared to the non-severe form. For example, DENV has been shown to increase the expression of sphingolipids (ceramide and sphingomyelin) in mosquito cells(5).Sphingolipids have recently been identified as an important contributor to vascular integrity and preventing endothelial cell hyper-permeability(5). The intracellular biosynthesis of ceramide takes place through two pathways namely, the *de novo* synthesis and salvage pathways (6) Several studies highlighted the role of two pathways in altering the life cycle and replication of some viruses (7–9). Thus, validating the role of selected lipids in disease progression during virus infection could shed light on understanding disease pathogenesis and develop novel therapeutics.Other studies have also shown that lipid changes could alter lipid microenvironments in the dengue disease(10).However, the association between lipid alterations and platelet count as well as haematocrit has not been studied. Therefore, this study aimed to determine the ceramide concentration in plasma from dengue patients using RP-HPLC and to assess their associations with the dengue severity symptoms.

## Methods

### Patients and study design

The study was approved by the Institutional Human Ethical Review Committee (ECR/451/Inst/PO/2013/RR-16), and all patients gave written informed consent. The patient presented to General Medicine at Mahatma Gandhi Medical College and Research Institute, Pondicherry with symptoms suggestive of dengue and confirmed by antigen (NS1) and an antibody test (IgM, IgG) were included in the study. Patients will be classified according to WHO 2009 criteria into two groups, namely, Group A, which includes severe dengue (SD-10) and non-severe dengue, which is further sub-categorised into dengue with warning signs (DWW-10) and dengue without warning signs (DWOW-10). Group B subjects (controls) are patients with other febrile illnesses (OFI-10). They would be defined as those patients who are negative for dengue NS1 antigen and anti-IgM/IgG dengue antibodies. About 3 mL of blood was collected in a tube containing 10% EDTA as an anticoagulant. Plasma was separated from the whole blood by centrifuging at 3000 g for 10 minutes and stored at -80°C.The ceramide (d18:1/22:0) levels from the plasma samples were thereafter measured.

### Lipid Extraction Protocol

### Reagents and Chemicals

HPLC grade methanol (Thermo Fisher Scientific (Sydney, Australia), 1-butanol (AR grade, Asia Pacific Specialty Chemicals (APS), Acetonitrile (Honeywell, Korea), Ammonium formate (Honeywell, Germany), Milli-Q water, were used for lipid extraction and quantification. Lipid standard - Ceramide (d18:1/22:0) (Avanti Polar Alabaster, Alabama, USA) was added to all test samples before lipid extraction (10 μL of a 1:10 dilution of Avanti polar lipids to 10 μL of neat plasma) for normalisation of raw peak areas and to correct for differences in extraction**(11)**.

### Quantification of ceramide in plasma

Lipids were extracted by the Alshehry Extraction Method (Single Phase 1-Butanol/Methanol 1:1 (v/v)) as previously described(12). The mobile phase was prepared by dissolving 0.0038 M of K2HPO4 in water and methanol, then sonicating. The organic phases were evaporated and the dried lipids were dissolved in a mixture (50 μl) of methanol and 0.07 M K2HPO4 (9:1 v/v). A derivatization mixture of 10 mg of o-phthaldialdehyde, 200 μl of ethanol, 10 μl of 2-mercaptoethanol, and 10 ml of boric acid (3% v/w) was prepared (pH 10.5). 5μl of the derivatization mixture, 45 μl of 9:1 ratio methanol, and 0.07 M K2HPO4 were added. The derivatives were analysed with a Shimadzu (Shlm-pack HPLC column system using an RP 18 Shimadzu column (4.6 mm × 250 mm) maintained at 40°C. Followed by, 20 μl of sample mixture being injected into the column. Then, set the flow rate at 1.000 mL/min for 20 minutes. The retention time of the selected peak was observed with the emission wavelength at 274 nm.

### *In vitro* study growth and maintenance using the HUVEC cell model

Human Umbilical Vein Endothelial Cells (HUVEC) purchased from Lonza (Walkersville, USA) were maintained in the laboratory as prescribed (13). In brief, HUVECs were seeded (1.5 ×10^4^/well) in a 6-well plate. Cells were starved in EBM2s basal medium for 24 hours at 37° C and 5% CO2 before incubation in media containing 10% human serum collected from SD, DWW, and OFI plasma samples(14). After 24 h of treatment, the HUVEC cells were harvested and resuspended in PBS solution. The levels of ceramide in the cells were estimated by HPLC methods.

### Statistical analysis

Results were expressed as median (interquartile range) or n (%). *P*-values of 0.05 or less were considered statistically significant. The normality test (Shapiro-Wilk Test) was performed to analyse the data. A comparison within the groups was performed using a two-tailed related sample t-test (Wilcoxon Signed Rank Test). Similarly, a comparison between the groups was performed using a two-tailed Independent Sample t-test (Mann-Whitney U test). The correlation was assessed using Spearman’s Rho correlation. A correlation analysis was directed to study the association between haematocrit and ceramide. All statistical tests were performed using SPSS software version 23 (SPSS, Chicago, IL, USA).

## Results

During the defervescence phase of dengue illness, platelet numbers fall as ceramide levels rise. In our current study, total plasma ceramide levels at the time of admission were elevated significantly (*P≤ 0.05) during the defervescence phase of infection, releasing an excess of ceramides into circulation.

### Plasma ceramide concentration

**Table 1.**
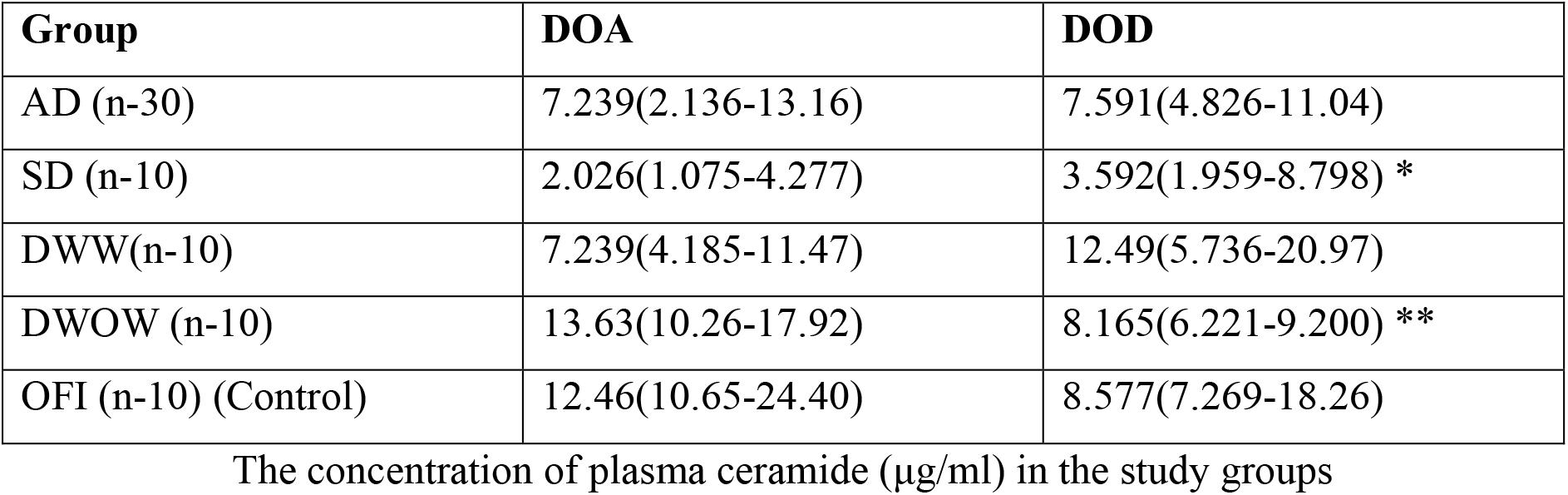
shows the levels of plasma ceramide in the study groups. Except for DWOW; there was a significant reduction in ceramide levels in SD, DWW, and AD cases as compared to OFI at the time of admission (Fig 1). Within the dengue groups, the severe group noted a significant (P≤ 0.01) drop in lipid levels as compared to the non-severe group.

| Group | DOA | DOD |
| --- | --- | --- |
| AD (n-30) | 7.239(2.136-13.16) | 7.591(4.826-11.04) |
| SD (n-10) | 2.026(1.075-4.277) | 3.592(1.959-8.798) * |
| DWW(n-10) | 7.239(4.185-11.47) | 12.49(5.736-20.97) |
| DWOW (n-10) | 13.63(10.26-17.92) | 8.165(6.221-9.200) ** |
| OFI (n-10) (Control) | 12.46(10.65-24.40) | 8.577(7.269-18.26) |

The concentration of plasma ceramide ( $\mu\text{g/ml}$ ) in the study groups

### Plasma levels of ceramide within the DOA and DOD

Furthermore, in dengue groups such as SD and DWOW patients, plasma levels of ceramides increased drastically towards defervescence (Fig.2). However, there was an enhanced amount of ceramide in DWW that had no significance (Fig 2). Ceramide levels in the dengue and OFI groups did not change significantly from the time of admission. Ceramide levels were found to be higher in the DWOW and AD groups during the defervescence phase of dengue infection.

**Figure 1.**
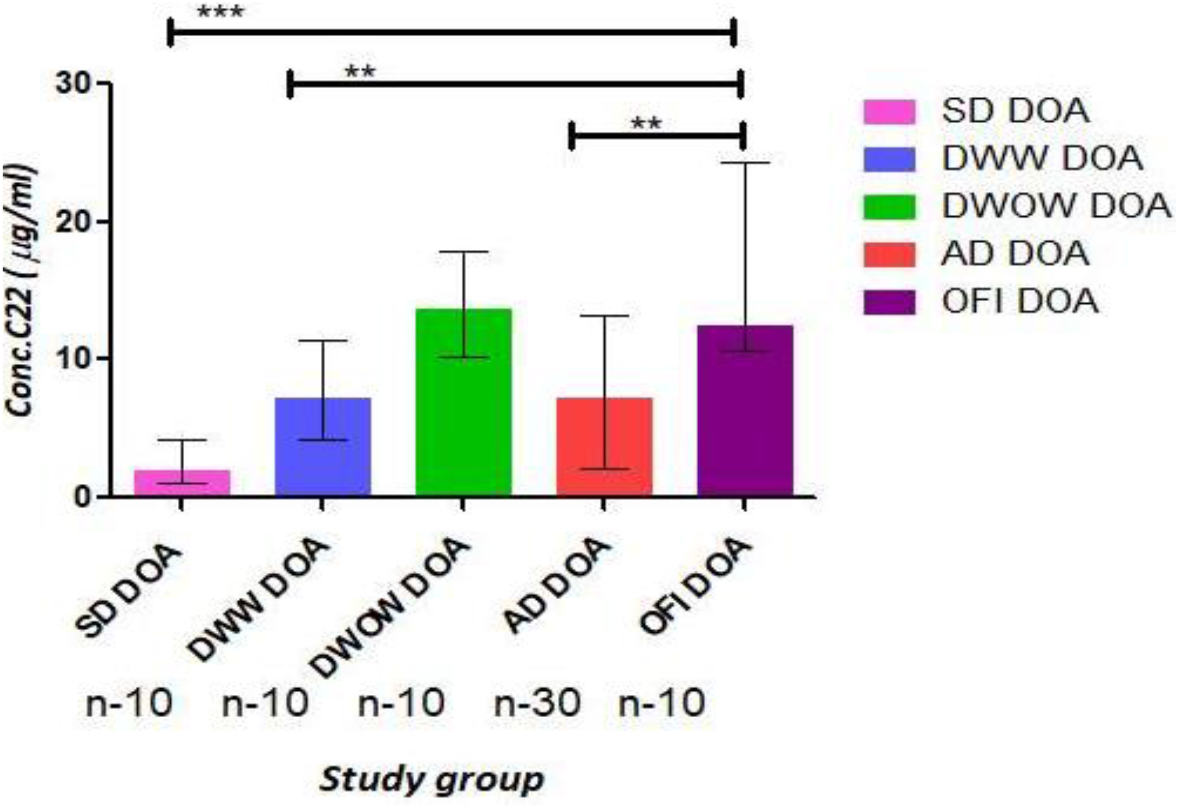
Decreased levels of ceramide (C22) at the time of admission. Statistical analysis was done using two-tailed Mann–Whitney tests to compare the data between the groups since the data was found to be non-parametric. P ≤ 0.05 was considered statistically significant. **AD** – All Dengue; **DWOW** - Dengue without Warning Sign; **DWW**-Dengue with Warning Sign, **SD** – Severe Dengue; **OFI** - Other Febrile Illness.

**Figure 2:**
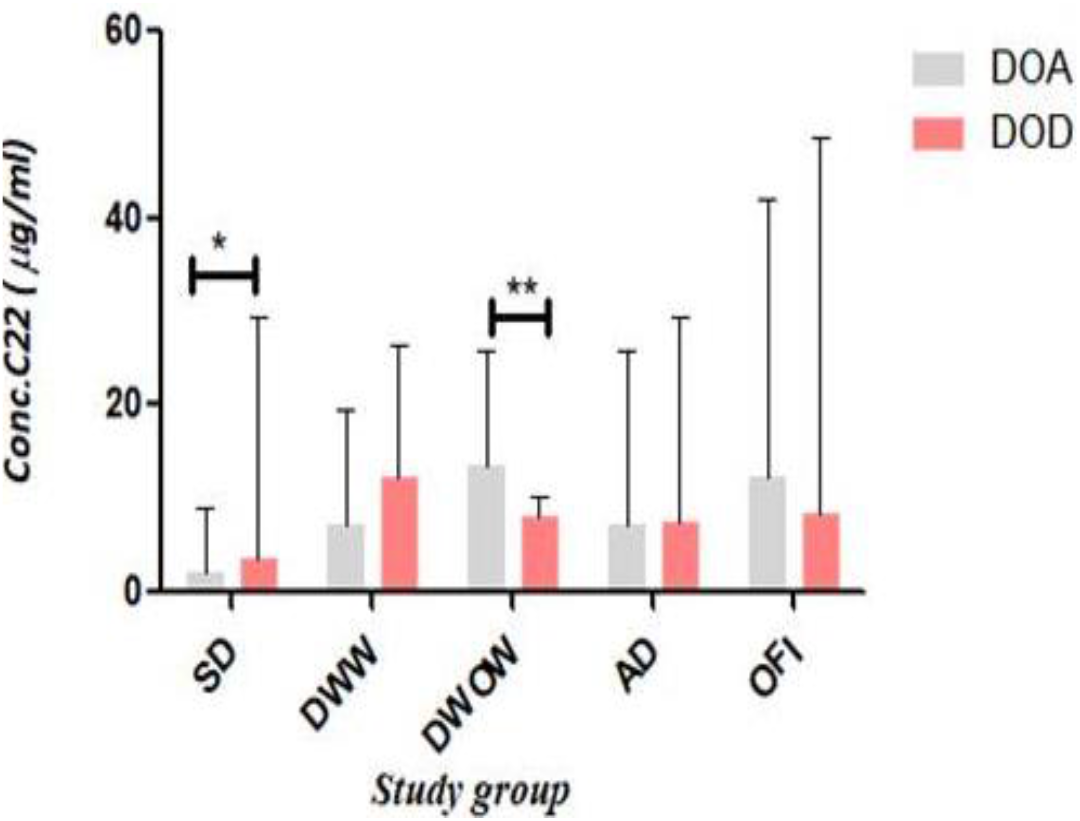
Ceramide levels are elevated in dengue patients. In comparison to OFI, our findings show that there is no significant difference across dengue groups in the febrile phase. Statistical analysis was done using two-tailed Wilcoxon signed-rank tests to compare data within groups since the data was found to be non-parametric. P≤0.05 was considered statistically significant. **DOA**-Day of admission and **DOD**-Day of defervescence

### Comparison between different groups at the time of admission and defervescence

When compared to control, there was a significant (P ≤ 0.05) downregulation of ceramides in the SD, DWW, and DWOW during the febrile phase (**Fig 3A**). In contrast, towards the critical phase, we noted a significant (P≤0.05) decrease in the levels of ceramide 22 in SD compared to DWW and OFI (**Fig 3B**).

**Figure 3:**
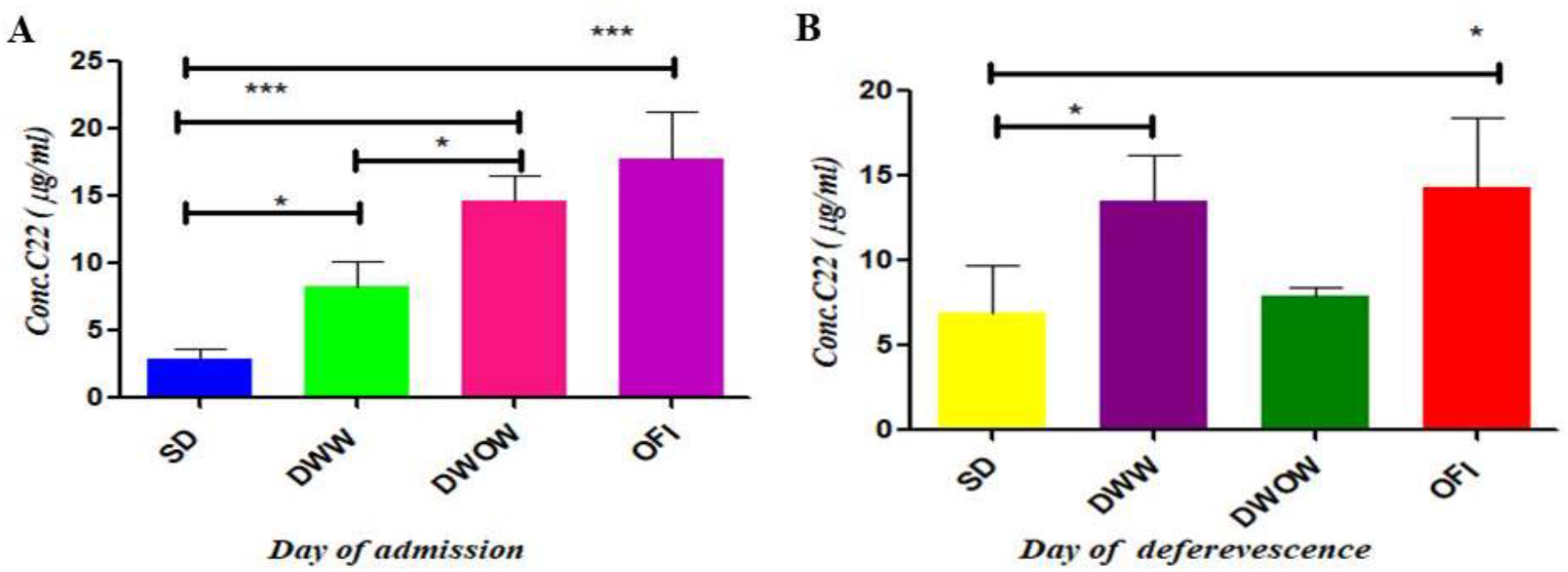
Ceramide concentrations in plasma (µg/ml) differed between the Dengue and OFI groups. In the Fig 3A and 3B, we depict the day of admission and Day of defervescence between the dengue groups and control. Statistical analysis was done using two-tailed Mann– Whitney tests to compare the data between the groups since the data was found to be non-parametric. P≤0.05 was considered statistically significant.

### Plasma levels of ceramides in HUVECs— *In vitro* study

The vascular endothelium serves as a barrier between the blood and the body’s surrounding tissue, which is a vital function. Ceramides may play a protective effect in improving vascular barrier function by reducing vascular leakage by maintaining tight junctions between endothelial cells. We examined the total plasma ceramide levels by RP-HPLC using the HUVECs model to look for changes in the endothelial cells.

### Association between haematocrit and ceramide levels

A negative correlation was observed between the plasma concentration of ceramide and haematocrit during the admission (R = -0.443, P≤ 0.05) and defervescence phase (R = -0.355, P≤0.05) in the AD group (Fig. 5A & B). However, no such correlation was observed in other dengue groups (SD, DWW, and DWOW) during infection. This result suggests that the lipid may be associated with vascular integrity, yet the role of this ceramide in endothelial leakage needs further investigation.

**Figure 4:**
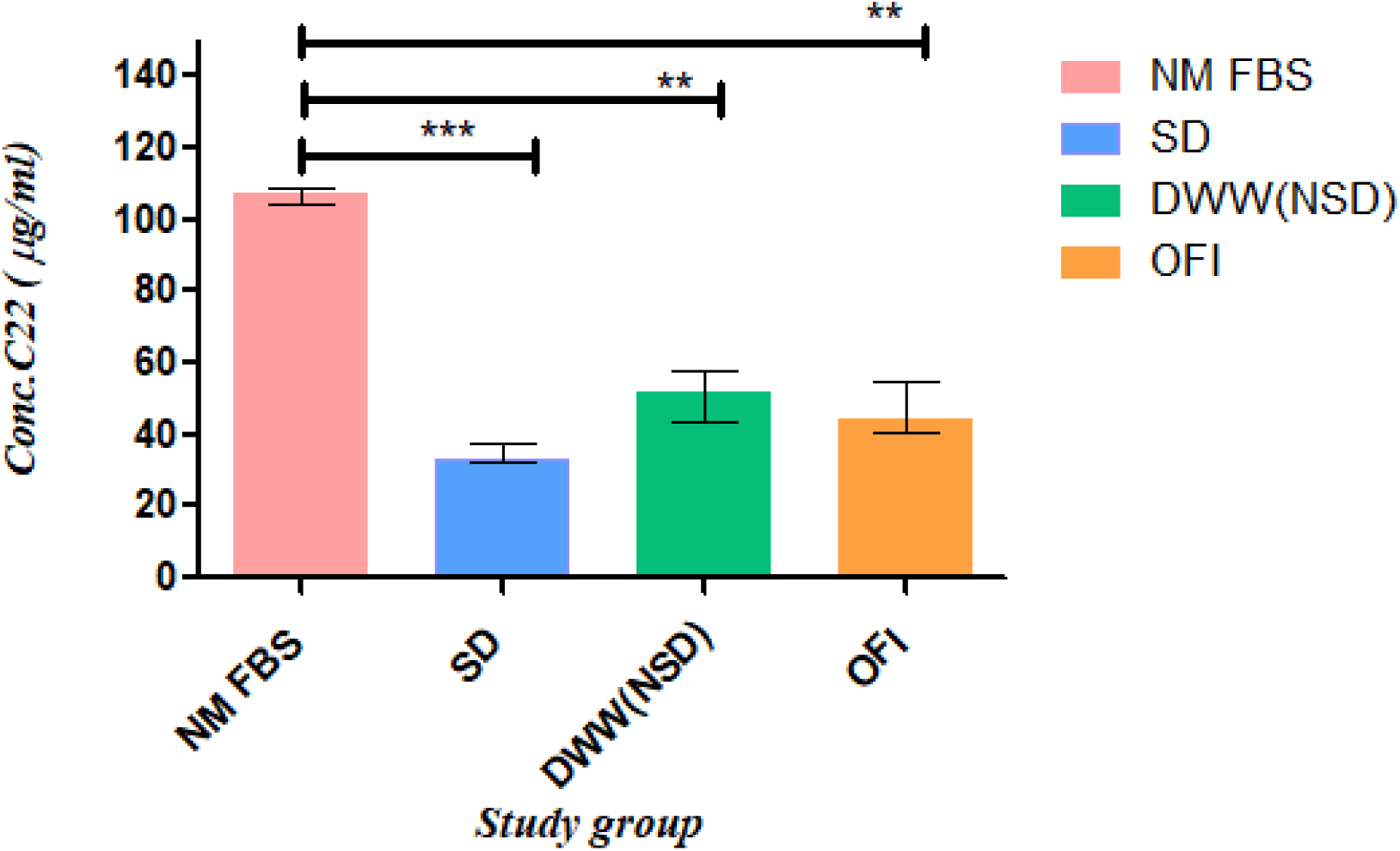
Comparison of Normal Foetal Bovine Serum (FBS) to other study groups. HUVEC cells treated with serum obtained from a severe form of dengue showed a significant decrease in the levels of ceramide compared to the cells treated with normal FBS. There were no significant differences observed when comparing the study groups and OFI. Median and interquartile ranges are shown. P≤ 0.05 was considered statistically significant.

**Figure 5:**
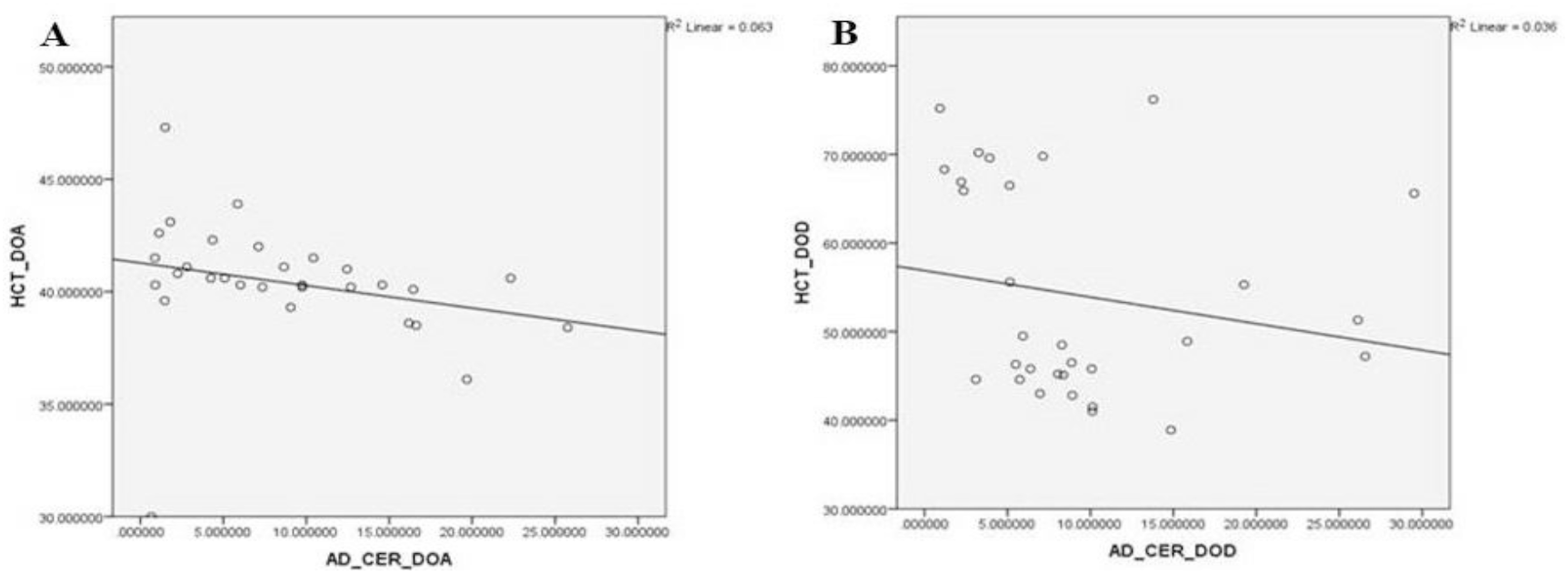
A negative correlation between ceramide and haematocrit during dengue. In Fig. 5 A and B, there is a correlation between ceramide and haematocrit during febrile phase (R = - 0.355, P≤0.05) and the defervescence phase (R = -0.443, P≤0.05) of the AD group. Since the data was nonparametric, Spearman’s Rho correlation was used to compare the data. A P-value ≤ 0.05 is considered significant.

### Relationship between Ceramide levels and Platelet count

We performed a correlation analysis to assess the relationship between platelet count and ceramide. A significant negative correlation was observed during the defervescence phase of the SD group (R =-0.867, P≤0.001) (Fig 6 A & B). But no such correlation was noted in other dengue groups (DWW, DWOW, and AD) throughout the infection.

**Figure 6:**
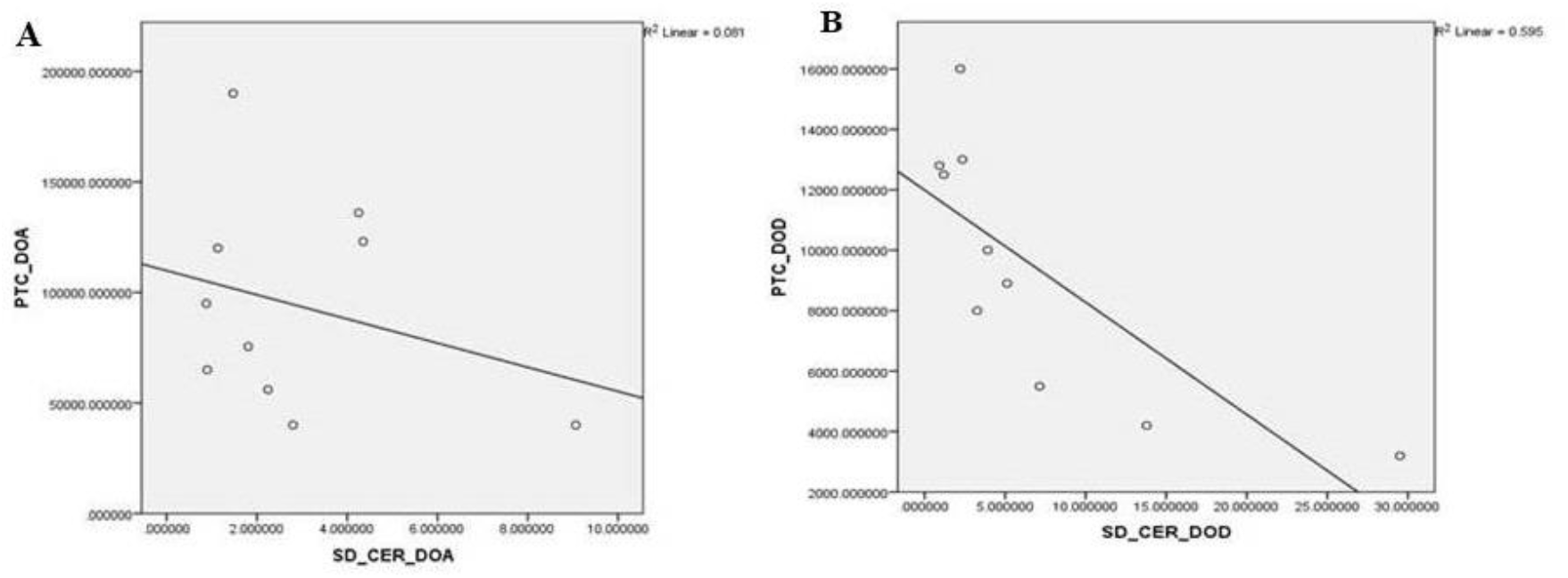
The negative correlation between ceramide and platelet counts during dengue. There was a correlation between ceramide and platelet count during the defervescence phase of the SD group (R = -0.867, P≤0.001). Since the data was nonparametric, Spearman’s Rho correlation was used to compare the data. A P-value ≤ 0.05 is considered significant.

## Discussion

A lipid in the pathogenesis of dengue is not explored much. We have reported a set of 23 differentially expressed platelet lipids in dengue patients(4).The present study assessed plasma levels of ceramide (d18:1/22:0) in dengue cases compared to other febrile controls. Previous studies suggested that ceramide and downstream products would interfere with the various stages of the life cycle of influenza A virus including replication, protein transport, and assembly(15-16). Ceramide is reported to have a role in the virus propagation like WNV, dengue(17).Tafesse et al. (2019) demonstrated that ceramide is a key signalling mediator in ZIKV replication and pathogenesis(18).Another study showed that treatment of cells with SMase markedly enhanced Japanese encephalitis virus infection and also indicates that ceramide plays a crucial role in the entry and exit of JEV into host cells(19).A significant decrease in the levels of ceramides was observed in SD, DWW, and AD cases compared to OFI at the time of admission, except for DWOW. The plasma level of ceramides was found to be decreasing towards DOD in DWOW cases, whereas the study observed a significant increase in ceramide levels towards DOD in SD and DWW cases. This enhanced ceramide found only in severe cases suggests the role of ceramides in the disease pathogenesis which typically lasts 24–48 hours during the defervescence phase. Most patients clinically improve during this phase, but those with substantial plasma leakage could develop severe dengue as a result of a sudden increase in vascular permeability. As a result, platelet count decreases, and ceramide levels eventually rise during infection. Together, these results show that ceramide is differentially expressed in severe dengue and patients characterised by severe symptoms during a critical phase. But it is still unclear whether the accumulation of ceramide in circulation could be attributed to vascular damage and platelet dysfunction, which could provide some light on the disease’s pathogenesis. However, an elevated level of ceramides in circulation may be a counter response by the host against virus infection.

The molecular basis by which ceramides impact endothelial cells remains poorly understood (20).The hyper-host responsive factors may be involved in the down regulation of ceramides during dengue infected patients. In support of this notion, the HUVEC cells treated with positive serum obtained from the critical phase of severe dengue showed a decrease in the levels of ceramide compared to the cells treated with normal FBS and non-severe dengue groups (DWW serum), serum from SD cases during febrile phase as well as OFI. This shows that the host inflammatory response can alter the ceramides in ECs during the course of dengue infection. In in vitro study, experiments indicated that Zika infection of microglial cells indicates up-regulation of ceramide and sphingosine in 48 hpi infection(21).

Plasma leakage is caused by increased capillary permeability and may manifest as haemoconcentration, as well as pleural effusion, hypoproteinaemia, and ascites(14).The 20% increase in haemoglobin levels(haematocrit) in severe dengue is a widely used sign of plasma leakage(22).In our study, a significant negative correlation (P≤ 0.05) was observed between the plasma concentration of ceramide and haematocrit during the time of admission and the defervescence phase of the AD group.

A study reported that sphingosine-1-phosphate enhances endothelial integrity while ceramide tends to promote vascular leak(23). Indeed, there is more evidence to support the elevated levels sphingolipids, including ceramides, exhibit an inhibitory effect during the entry of viral infection. Besides viral infections, plasma ceramide levels are elevated in inflammatory conditions such as hypertension, diabetes, and cardiovascular diseases(24-25).The differentially expressed plasma lipids levels increase in SD reveals its importance in disease pathogenesis. Though, plasma levels of ceramides showed a negative association with HCT as well platelet count, the exact role of ceramides in endothelial dysregulation is not known. Perhaps, further investigations are required to ascertain the ceramides as an effective prognostic marker in a large clinical setup.

## Conclusion

A negative association between ceramide and platelet count could be used as early indicator for disease progression. An increased levels of ceramide (d18:1/22:0) towards the defervescence phase during dengue infection may serve has a robust biomarker for the early prediction of disease severity.

## Abbreviation

SD: Severe dengue
DWOW: Dengue without warning sign
DWW: Dengue with warning sign
AD: All dengue
OFI: Other febrile illnesses
HPLC: High-performance liquid chromatography
DOA: Day of Admission
DOD: Day of Defervescence
HUVECs: Human umbilical vein endothelial cells
ECs: Endothelial cells
FBS: Foetal bovine serum
HCT: Haematocrit
WNV: West Nile virus
DENV: Dengue virus
HCV: Hepatitis C virus
HIV: Human Immunodeficiency virus
JEV: Japanese encephalitis virus
ZIKV: Zika virus

## Acknowledgment

The authors also thank the medical staff of the MGMCRI for the collection of the blood samples. We also acknowledge the infrastructure support provided by Sri Balaji Vidyapeeth for conducting this research and writing this manuscript.

## Ethics

The study protocol has been approved by the Institutional Human Ethics committee (ECR/451/Inst/PO/2013/RR-16)

## Reference

1. Harapan H, Ryan M, Yohan B, Abidin RS, Nainu F, Rakib A, et al. Covid-19 and dengue: Double punches for dengue-endemic countries in Asia. Rev Med Virol. 2020 Sep 18;e2161.

2. Matsunaga K ichiro, Kimoto M, Lim VW, Thein TL, Vasoo S, Leo YS, et al. Competitive ELISA for a serologic test to detect dengue serotype-specific anti-NS1 IgGs using high-affinity UB-DNA aptamers. Sci Rep. 2021 Sep 9;11(1):18000.

3. Balakrishna Pillai AK, Chu JJH, Mariappan V, JeanPierre AR. Platelets in the pathogenesis of flavivirus disease. Current Opinion in Virology. 2022 Feb 1;52:220–8.

4. Samadanam DM, Muthuraman KR, Mariappan V, Kadhiravan T, Parameswaran N, Balakrishna Pillai AK, et al. Altered Platelet Fatty Acids in Dengue Cases by Gas Chromatography-Mass Spectrometry Analysis. INT. 2019;62(2):57–64.

5. Yager EJ, Konan KV. Sphingolipids as Potential Therapeutic Targets against Enveloped Human RNA Viruses. Viruses. 2019 Oct 1;11(10):912.

6. Castro BM, Prieto M, Silva LC. Ceramide: a simple sphingolipid with unique biophysical properties. Prog Lipid Res. 2014 Apr;54:53–67.

7. Dai L, Trillo-Tinoco J, Bai A, Chen Y, Bielawski J, Del Valle L, et al. Ceramides promote apoptosis for virus-infected lymphoma cells through induction of ceramide synthases and viral lytic gene expression. Oncotarget. 2015 Jul 3;6(27):24246–60.

8. Kanj SS, Dandashi N, El-Hed A, Harik H, Maalouf M, Kozhaya L, et al. Ceramide regulates SR protein phosphorylation during adenoviral infection. Virology. 2006 Feb 5;345(1):280–9.

9. Sphingosine Kinase-2 Maintains Viral Latency and Survival for KSHV-Infected Endothelial Cells [Internet]. [cited 2022 Apr 22]. Available from: https://journals.plos.org/plosone/article?id=10.1371/journal.pone.0102314

10. Heaton NS, Randall G. Multifaceted roles for lipids in viral infection. Trends Microbiol. 2011 Jul;19(7):368–75.

11. Wong MWK, Braidy N, Pickford R, Sachdev PS, Poljak A. Comparison of Single Phase and Biphasic Extraction Protocols for Lipidomic Studies Using Human Plasma. Front Neurol. 2019 Aug 21; 10:879.

12. Matyash V, Liebisch G, Kurzchalia TV, Shevchenko A, Schwudke D. Lipid extraction by methyl-tert-butyl ether for high-throughput lipidomics. J Lipid Res. 2008 May;49(5):1137–46.

13. Cardozo FTG de S, Baimukanova G, Lanteri MC, Keating SM, Ferreira FM, Heitman J, et al. Serum from dengue virus-infected patients with and without plasma leakage differentially affects endothelial cells barrier function in vitro. PLOS ONE. 2017 Jun 6;12(6):e0178820.

14. Mariappan V, Adikari S, Shanmugam L, Easow JM, Balakrishna Pillai A. Expression dynamics of vascular endothelial markers: endoglin and syndecan-1 in predicting dengue disease outcome. Transl Res. 2021 Jun; 232:121–41.

15. McVey MJ, Weidenfeld S, Maishan M, Spring C, Kim M, Tabuchi A, et al. Platelet extracellular vesicles mediate transfusion-related acute lung injury by imbalancing the sphingolipid rheostat. Blood. 2021 Feb 4;137(5):690–701.

16. Seo YJ, Blake C, Alexander S, Hahm B. Sphingosine 1-Phosphate-Metabolizing Enzymes Control Influenza Virus Propagation and Viral Cytopathogenicity. Journal of Virology. 2010 Aug 15;84(16):8124–31.

17. Aktepe TE, Pham H, Mackenzie JM. Differential utilisation of ceramide during replication of the flaviviruses West Nile and dengue virus. Virology. 2015 Oct 1; 484:241–50.

18. Leier HC, Messer WB, Tafesse FG. Lipids and pathogenic flaviviruses: An intimate union. PLoS Pathog. 2018 May 10;14(5): e1006952.

19. Tani H, Shiokawa M, Kaname Y, Kambara H, Mori Y, Abe T, et al. Involvement of Ceramide in the Propagation of Japanese Encephalitis Virus. J Virol. 2010 Mar 15;84(6):2798–807.

20. Srikiatkhachorn A. Plasma Leakage in Dengue Hemorrhagic Fever. Thromb Haemost. 2009 Dec;102(6):1042–9.

21. Diop F, Vial T, Ferraris P, Wichit S, Bengue M, Hamel R, et al. Zika virus infection modulates the metabolomic profile of microglial cells. PLoS One. 2018 Oct 25;13(10): e0206093.

22. Ralapanawa U, Alawattegama ATM, Gunrathne M, Tennakoon S, Kularatne SAM, Jayalath T. Value of peripheral blood count for dengue severity prediction. BMC Res Notes. 2018 Jun 20; 11:400.

23. Jernigan PL, Makley AT, Hoehn RS, Edwards MJ, Pritts TA. The role of sphingolipids in endothelial barrier function. Biol Chem. 2015 Jun;396(6–7):681–91.

24. Spijkers LJA, van den Akker RFP, Janssen BJA, Debets JJ, De Mey JGR, Stroes ESG, et al. Hypertension Is Associated with Marked Alterations in Sphingolipid Biology: A Potential Role for Ceramide. Najbauer J, editor. PLoS ONE. 2011 Jul 19;6(7): e21817.

25. Predescu S, Knezevic I, Bardita C, Neamu RF, Brovcovych V, Predescu D. Platelet Activating Factor-Induced Ceramide Micro-Domains Drive Endothelial NOS Activation and Contribute to Barrier Dysfunction. Zhao YY, editor. PLoS ONE. 2013 Sep 27;8(9): e75846.

